# A re-examination of responding on ratio and regulated-probability interval schedules

**DOI:** 10.1101/284216

**Authors:** Omar D. Pérez, Michael R.F. Aitken, Amy L. Milton, Anthony Dickinson

**Author notes:** Corresponding author: Omar D. Pérez, Division of Humanities and Social Sciences, California Institute of Technology, Pasadena, 1200 East California Boulevard, California, CA 91107, USA.

## Abstract

The higher response rates observed on ratio than on matched interval reward schedules has been attributed to the differential reinforcement of longer inter-response times (IRTs) on the interval contingency. Some data, however, seem to contradict this hypothesis, showing that the difference is still observed when the role of IRT reinforcement is neutralized by using a regulated-probability interval schedule (RPI). Given the mixed evidence for these predictions, we re-examined this hypothesis by training three groups of rats to lever press under ratio, interval and RPI schedules across two phases while matching reward rates within triads. At the end of the first phase, the master ratio and RPI groups responded at similar rates. In the second phase, an interval group yoked to the same master ratio group of the first phase responded at a lower rate than the RPI group. Post-hoc analysis showed comparable reward rates for master and yoked schedules. The experienced response-outcome rate correlations were likewise similar, and approached zero as training progressed. We discuss these results in terms of dual-system theories of instrumental conditioning.

## Introduction

Two basic patterns of reward delivery are commonly used in instrumental conditioning experiments. In the first class, *ratio* schedules, reward delivery depends only on the number of responses performed, so that a reward is delivered every time a response requirement is attained. In the second class, *interval* schedules, the delivery of rewards depends not only on responding but also on time; a reward is scheduled to be delivered after an elapsed period of time, such that the first response performed after this time period is rewarded. The random-ratio (RR) and random-interval (RI) schedules are idealized stochastic versions of these schedules. The RR schedule sets a fixed reward probability per response, say *q*, so that the (1/ *q*)*th* response is, on average, rewarded; the RI schedule programs the availability of a reward with a fixed probability, say *r*, in each second: rewards are therefore collected with probability 1 after an average of 1/ *r* sec have elapsed.

An enduring debate in instrumental learning concerns the mechanisms driving performance under ratio and interval training. For the most part, learning theorists have argued that reward probability is the primary determinant of instrumental performance because response rates increase as the reward probability per response increases. However, the fact that ratio and interval performance differs even when the reward probability is matched (Catania et al., 1977; Dickinson, Nicholas, & Adams, 1983; Peele, Casey, & Silberberg, 1984; Reed, 2001; Skinner, 2014; Zuriff, 1970) suggests that additional variables are involved in the instrumental learning process. Moreover, the ratio-interval difference persists when the reward rate (rather than reward probability) is matched, such that the reward probability experienced by interval-trained subjects is higher than the one experienced by those trained under the ratio schedule. This widely-observed result is problematic for theories of instrumental learning based on reward probability, both associative (Mackintosh, 1974; Mackintosh & Dickinson, 1979) and computational (Daw, Niv, & Dayan, 2005; Dezfouli & Balleine, 2012; Keramati, Dezfouli, & Piray, 2011; Niv, Daw, & Dayan, 2005).

The question at issue is what produces this difference. If the incentive value and the probability or rate of reward are the same, the expected value of both training regimes should be equal. So why does ratio training elicit more effort? To answer this question, probability-based theories shift the focus from individual responses to the pause between responses, or inter-response time (IRTs). On interval schedules, it is argued, longer IRTs yield higher reward probabilities for the next response that is to be emitted (Dawson & Dickinson, 1990; Peele et al., 1984; Tanno & Silberberg, 2012; Wearden & Clark, 1988). If the animal is sensitive to the differential reinforcement of longer IRTs, interval responding should hence be slowed compared to a ratio schedule under which reinforcement probability remains constant with variations in IRT. Even though the reward probability *per response* programmed by the experimenter might be similar, these theories stress the importance of the *joint* probability per response together with the time elapsed since the last reward to explain responding (Niv, Daw, Joel, & Dayan, 2007; Peele et al., 1984; Shimp, 1973; Wearden & Clark, 1988).

One way of investigating the sufficiency of IRT-reinforcement in explaining instrumental performance is to design a schedule in which this factor is neutralized while keeping the average interval between rewards constant at the scheduled value. The regulated-probability interval schedule (RPI), originally designed by Kuch and Platt (Kuch & Platt, 1976), does this by continuously recording the local response rate and then setting the reward probability for the next response (*P*) to a value that maintains the scheduled reward rate if the animal continues to respond at the same rate. Thus, the reward probability for the next response is independent of the preceding IRT. Formally, if *T* is the scheduled interval between rewards, the RPI sets the reward probability for the *next* performed response to *P* = *t* / *Tm*, where *t* is the time it has taken the subject to perform the last *m* IRTs. Thus, in the RPI the reward probability is not determined entirely upon the current IRT, but on the duration of a number (*m*) of IRTs emitted before the current IRT. The RPI, in other words, considers a *local* response rate *B*_*m*_ = (*m* +1) / *t* given by the last m+1 responses (or *m* IRTs) in the last *t* secs, and fluctuates the reward probability inversely with respect to this local response rate so that the agent still receives a constant reward rate of 1/ *T* rewards per second independently of the length of the current IRT. Since the current magnitude of IRT contributes only a fraction 1/ *m* of the change in reward probability, the RPI should also be able to neutralize the effect of timing on increasing reward probabilities for long IRTs.

If IRT-reinforcement is sufficient to explain the difference between ratio and interval responding, response rates under RPI schedules should be higher than under RI training. In addition, since IRT size is independent of reward probability in both ratio and RPI schedules, responding should be comparable between these two schedules.

The evidence for these predictions is, however, both mixed and scarce. In a within-subject study of lever pressing, Tanno and Sakagami (2008) trained four rats under ratio, RPI and interval schedules and found that ratio and RPI training maintained comparable response rates. Consistent with the IRT-reinforcement hypothesis, they also observed lower responding under interval schedules than under both ratio and RPI schedules. However, in a previous study of chain pulling, Dawson and Dickinson (1990) trained three triads of rats with respect to reward rate and found higher responding on ratio than RPI schedules, suggesting that IRT-reinforcement cannot be the only variable involved in the ratio-interval difference. Furthermore, the fact that theories of instrumental conditioning have been mostly informed by lever-pressing data makes it important to further examine this hypothesis using this target response. This was the goal of the present experiment.

The present study comprised two phases of training and three groups of rats (see 1). In the first phase, the programmed reward rate for a RPI group was yoked to that generated by a master RR group. As IRT-reinforcement theory anticipates no difference between the performance of RR and RPI groups for matched reward rates, it was necessary to show that performance was also sensitive to reward probability. To this end, a second RR group received rewards with a higher probability than the master RR group and therefore should have responded at a higher rate. In the second phase, this high-probability RR group was switched to a standard RI schedule with the reward rate yoked to the same master RR group of the first phase. The contrast between RPI and RI performance in this second phase established whether responding was sensitive to the differential reinforcement of long IRTs by yielding a lower response rate in the RI group.

## Method

### Subjects

The subjects were 36 male naïve Lister Hooded rats (Charles River, Margate, UK) that were around 3 months old and with a mean free-feeding weight of 374 g at the beginning of the experiment. They were caged in groups of four in a vivarium under a 12-hour reversed light-dark cycle (lights off at 0700). All rats had *ad libitum* access to water in their home cages. All rats were mildly food restricted throughout training by being fed for 1 hour in their home cages after every session. This research was regulated under the Animals (Scientific Procedures) Act 1986 Amendment regulations 2012 under Project Licence 70/7548 following ethical review by the University of Cambridge Animal Welfare and Ethical Review Body (AWERB).

### Apparatus

Subjects were trained in twelve operant chambers (Med Associates, Vermont, USA) controlled by Whisker Server (Cardinal & Aitken, 2009). A client written in Visual Basic © and run on a laptop computer (ASUS © K52J) under Microsoft Windows © 10 was utilized to communicate with the server, control the chambers and retrieve the data.

Each chamber had a magazine and two retractable levers at each side of the magazine. A pellet dispenser delivered 45-mg chocolate-flavored pellets (Sandown Scientific, Middlesex, UK) into the magazine. A 2.8-W house light illuminated the chamber during each experimental session.

### Behavioural procedures

#### Pretraining

Rats were first given two magazine training sessions with both levers retracted. During these sessions, the pellets were delivered on a random time 60-sec schedule (i.e. a reward was programmed to be delivered on average after 60 sec in the absence of the lever) until 30 rewards were delivered and consumed. On the next session one of the levers was inserted at the start of the sessions and each press was reinforced on a fixed ratio (FR1) schedule. Finally, for each of the three subsequent days, lever pressing was rewarded under an RR schedule with increasing ratio requirements until 30 rewards were earned. The parameter was 5 on Day 1, and 10 on Days 2 and 3. The active lever was counterbalanced between subjects, but each subject was trained with the same lever and in the same operant box throughout training. All rats were given two runs of pre-training until all rewards were delivered and consumed.

#### Training

On the following day subjects were randomly assigned to 3 different groups (N=12 each) (see Table 1) the first phase of the experiment, master rats in Group A were run under an RR-20 schedule, so that 1 of every 20 responses on average was rewarded. On a second run, yoked subjects in Group B were trained on a RPI schedule (RPI-y) with the same inter-reinforcement intervals produced by their corresponding master subjects in the RR-20 group. On a third run, group C was run under a RR-10 schedule, so that 1 of every 10 responses was rewarded.

**Table 1.** Design of the Experiment. The master group is written in bold. The subscript “y” signifies that the group was yoked to the master group with respect to reward rate. Each group was run for 10 sessions in each phase of the experiment. Responding was extinguished between each phase by giving rats 3 sessions of reward delivery non-contingently to lever pressing.

| Phase | Group |  |  |
| --- | --- | --- | --- |
|  | A<br>(N=12) | B<br>(N=12) | C<br>(N=12) |
| 1 (10 sessions) | **RR-20** | RPI-y | RR-10 |
| 2 (10 sessions) | **RR-20** | RPI-y | RI-y |

In the second phase of the experiment, rats in Group C were switched to a RI schedule yoked to the master rats in the RR-20 group in the same manner as rats in the RPI-y group (Group B), thereby yielding two interval groups yoked by reward rate within triads. Across all sessions of the experiment, the yoked rats were trained in the same operant chamber and with the same lever as their corresponding master rats. In order to minimize any carry-over effects between phases - particularly in Group C which was shifted from a ratio to an interval schedule – between the two phases all rats received three sessions during which the food pellets were delivered non-contingently to lever-pressing on a random-time 60-sec schedule.

To prevent the development of extreme patterns of response bursting (Reed, 2011; Shull, Gaynor, & Grimes, 2001), for all schedules tested in the experiment the reward schedule was programmed so that only the first response in each 1-sec window interrogated the reward probability in the computer; all other responses during these windows were recorded but did not engage the probability generator. Additionally, to prevent rats from stopping responding when a reward had taken too many responses to be earned or too long an interval to be scheduled, we constrained these parameters to a maximum of three times the nominal response or interval requirement, respectively. For example, if the number of responses currently performed since the last reward in the RR-20 schedule was 60 responses, the next response was rewarded. Likewise, in the RPI or RI schedules, if the current interval since last reward was 3*T*, and *T* is the scheduled interval, then the next response was rewarded. Since Dawson and Dickinson (1990) found no evidence of memory size in the RPI schedule affecting performance, we set the value of *m* to 50. The same memory size was employed by Tanno and Sakagami (2008) and Dawson and Dickinson (1990) in their previous work.

Since the variance in response rates increases with the mean, a square root transformation to this variable was applied for all statistical analysis. Welch t-tests and Cohen’s D (with 95% confidence interval) were calculated for the pre-planned contrasts of interest. The significance of the contrasts was evaluated against the standard criterion of *α* = .05. When no statistical difference was found between groups, a Bayes Factor (*BF*_01_) in favor of the null was also calculated to test the likelihood of the null over the alternative hypothesis of there being a difference between the groups in terms of the variable being analyzed (Morey & Rouder, 2015). All the analyses were performed using the R programming language running in RStudio (RStudio Team, 2015) and extended with the packages BayesFactor, reshape2 v.1.4.1, plyr v.1.8.3, and ggplot2 v.2.1.0. Data and scripts for all the analyses can be found at www.github.com/omadav/RPIrats.

## Results

### Phase 1: an RR10 schedule produced greater responding than an RR20 schedule, with equivalent responding in the RPI-y group

Figure 1A presents the response rates in the last three sessions of training for each group and phase of the study; the acquisition curves across all sessions are presented in Figure 1B. The results from Phase 1 supported the prediction of all instrumental theories with regard to RR schedules with different reward probabilities in that rats in the RR-10 group responded more vigorously than those in the RR-20 group, *t*(69.9) = −5.82, *p* < .01, *D* = −1.37 95% *CI* [−1.90, −0.85]. In contrast, there was no detectable difference in responding for the RR 20 and the RPI-y groups at the end of the first phase of training, *t*(68.7) = −1.39, *p* = .17, *D* = −0.33 95% *CI* [−0.80, 0.14], suggesting that R-O rate correlation did not have any detectable effect on increasing responding in the RR group.

**Figure 1.**
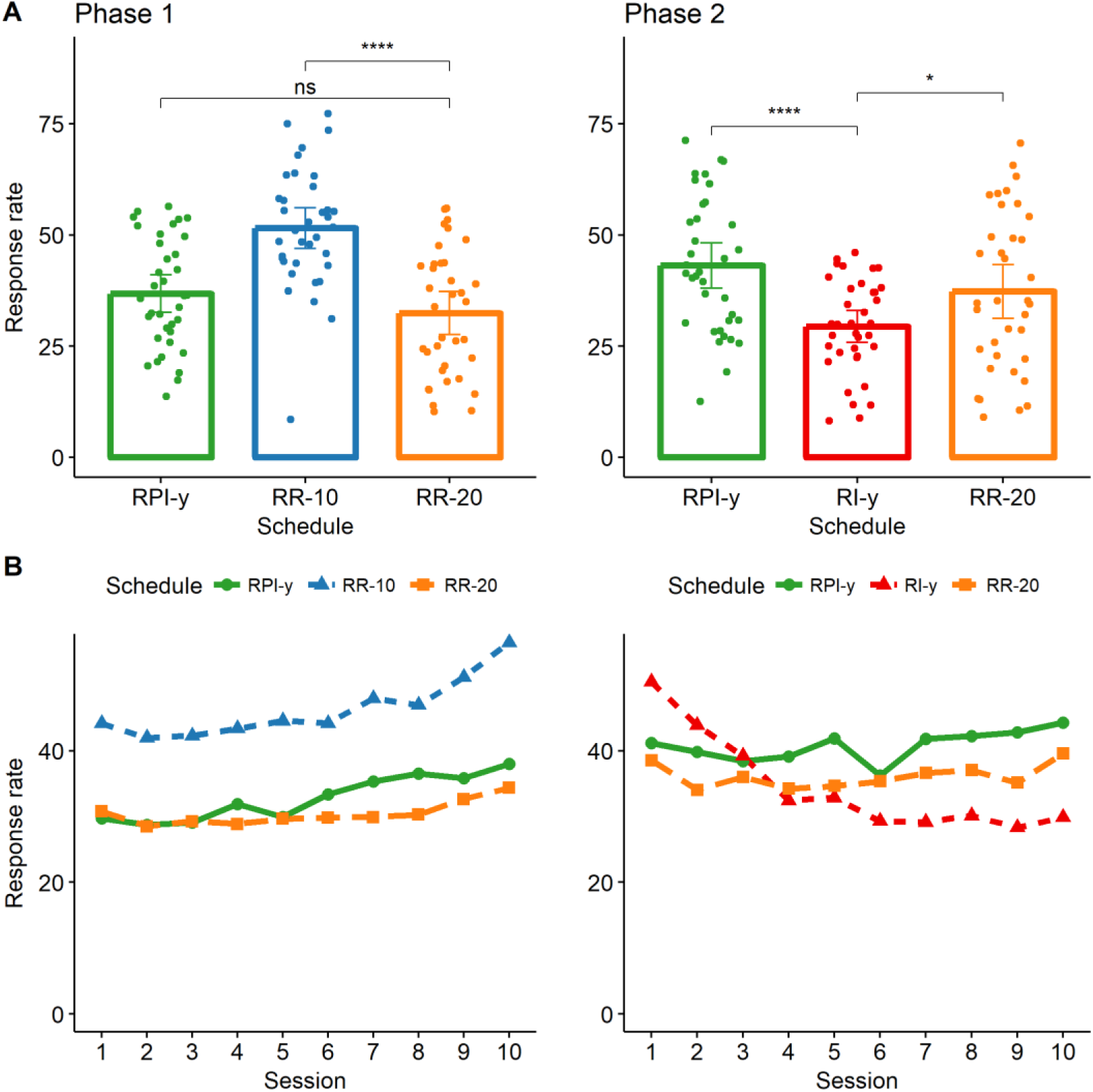
Response rates for phases 1 and 2 of the experiment. **A**. Average response rates maintained by rats in the last 3 sessions of training in each phase of the study. **B**. Acquisition curves for the 10 sessions of training in each phase of the study. * p<.05, ****p<.001. Error bars represent 95% bootstrapped confidence intervals.

### Phase 2: responding of an RI-y group was lower than both responding on an RR20 schedule and responding on an RPI-y schedule

To test the effect of differential IRT reinforcement on responding, in Phase 2 Group C (the RR-10 group from the previous phase) was switched to an RI-y schedule yoked to the master RR-20 group with respect to reward rate in the same way as the RPI-y group was yoked to the RR-20 group in both phases. The RR-20 group responded at a higher rate than the RI-y group, *t*(62.8) = 4.44, *p* < .01, *D* = 0.24 95% *CI*[−0.23, 0.72], replicating the widely-observed ratio-interval difference when regular interval schedules are employed (Dickinson et al., 1983; Peele et al., 1984) Importantly, the RI-y group also responded at a lower rate than the RPI-y group, *t*(57.1) = 2.27, *p* = .03, *D* = 0.57 95% *CI*[0.09,1.05], showing that the differential reinforcement for long IRTs was able to slow responding in the RI-y group compared to a reward that does not hold this property.

### Yoking analysis

The yoking analysis was performed separately for the last 3 sessions of each phase of the study. For each phase, possible differences between those variables that were intended to be matched by the yoking procedure were analyzed; in particular, the interest was put in ensuring the yoking was successful in matching reward rates and, in addition, whether the RPI schedule was successful in controlling the differential reinforcement of long IRTs. Table 2 shows the results for each phase of the experiment.

**Table 2.**
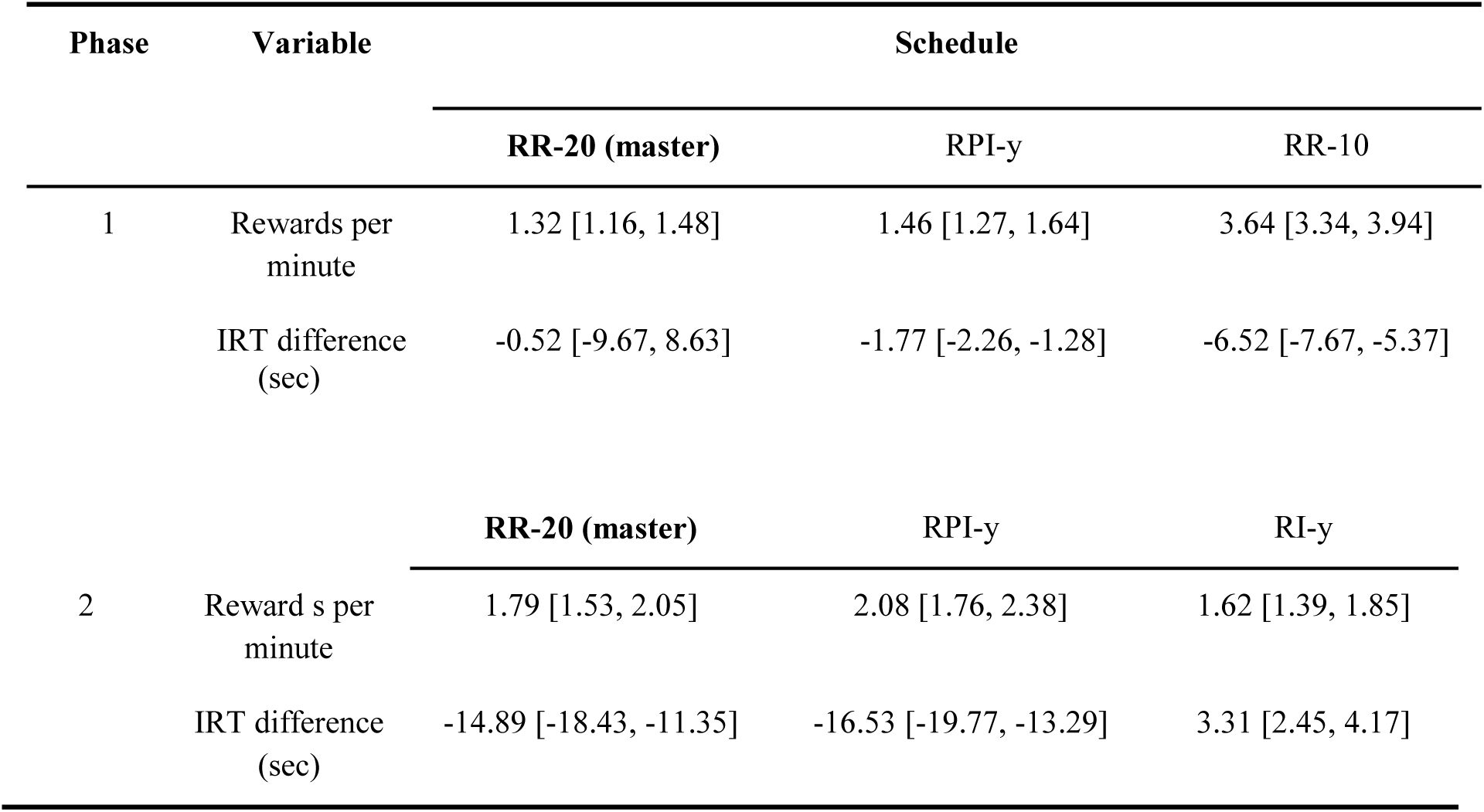
Results of the yoking procedure. Mean and 95% bootstrapped confidence intervals for Phase 1 and Phase 2 of the study. The reward rates are in rewards per min; the IRT difference is calculated as the difference between the mean reinforced IRT and the mean IRT emitted by each rat. The letter “y” signifies that the group was yoked to the reward rate obtained in the master ratio group. This was achieved by taking triads of rats, one in the master RR-20 group and one in each of the RPI-y and RI-y groups, and using the reward rate experienced by the master rat in each session as the parameter for the latter two rats in each session.

### Reward rate

The reward rates obtained in each session were similarly analyzed for the last 3 sessions of the two phases of the experiment (**Table 2**). The results confirmed that the yoking procedure succeeded in matching the reward rates with respect to the master RR 20 group (Phase 1: RR-20 - RPI-y:

*t*(68.8) = −1.15, *p* = .25, *D* = −0.27 95% *CI* [−0.74, 0.20], *BF*_01_ = 2.34; Phase 2: RR-20 - RPI-y:

*t*(67.9) = −1.42, *p* = .16, *D* = −0.33 95% *CI* [−0.81, 0.14], *BF*_01_ = 1.74; RR-20 - RI-y:

*t*(68.8) = 1.03, *p* = .30, *D* = 0.24 95% *CI* [−0.23, 0.72], *BF*_01_ = 2.60.)

### IRT reinforcement

Unlike RI schedules, the RPI schedule aims not to assign responses followed by longer IRTs a higher reward probability. The same property holds for the RR schedule, although for a simpler reason: in this case, the probability of obtaining a reward depends only on each response independently, and not on the time that has elapsed since the last reward obtained. Therefore, the difference between the mean rewarded IRT size and mean emitted IRT size is an index of whether the reward schedule was rewarding responses followed by long IRTs over all other emitted IRTs. The index, therefore, should be less than or equal to zero in RR and RPI schedules. In contrast, since the reward probability is a direct function of IRT size on the RI schedule, the value of this index for RI schedules should be positive. If both predictions are confirmed from the data, then the differences between the RPI and RI groups can be attributable to the latter schedule rewarding long pauses between responses in a higher proportion. The data (**Table 2**) confirmed that this was the case, as only the RI-y group had a positive index.

## Discussion

Using lever pressing in rats, the present study examined the predictions of IRT-reinforcement theories regarding RR, RI, and RPI performance under matched reward rates. The first phase confirmed the critical role of reward probability in instrumental responding, as indicated by the higher response rates observed for RR-10 compared to the RR-20 group. By contrast, there was no detectable difference in responding between the RR-20 group and the yoked RPI-y group. The second phase of the experiment confirmed the widely observed ratio-interval difference, in that the RR-20 group came to perform at a higher rate than the yoked RI-y group (Cole, 1994, 1999; Peele et al., 1984; Zuriff, 1970) at the end of training. Direct evidence for the role of differential reinforcement of long IRTs was revealed in the lower responding on the RI-y group compared to the RPI-y group in this second phase.

These data are in agreement with the results obtained by Tanno and Sakagami (2008), who found similar lever-pressing performance in RR and RPI schedules and higher responding on these two schedules compared to a RI schedule using a within-subject design. Although formal models have not presented explicit simulations for the RR-RPI contrast, these data are clearly amenable to both associative (Peele et al., 1984; Tanno & Silberberg, 2012; Wearden & Clark, 1988) and normative (Niv, 2007; Niv et al., 2007) models based on IRT reinforcement with regard to the ratio-interval comparison. By contrast, our data are at variance with those reported by Dawson and Dickinson (1990), who found higher chain-pulling performance on RPI compared to RI schedules, and lower performance on RPI than RR schedules yoking reward rates within triads.

Dawson and Dickinson (1990; see also Pérez et al., 2016; Dickinson, 1985) explained the higher response rate observed under ratio than RPI training by noting the different linear relationship between responding and rewards or outcomes established by the two schedules. This idea, originally espoused by Baum (1973) as an alternative to the probability-based Law of Effect (Thorndike, 1911), shifts the focus from single or discrete events to numerous events extended in time, called activities (Baum, 2002).^1^ The critical assumption is that agents will be able to encode the rate of responding and reward over an extended period and deploy this information to act in accordance with this relationship. Given the independence of reward probability and time established by ratio schedules, the reward rate will increase (decrease) every time the response rate increases (decreases) under this type of training; since on interval schedules the reward rate is fixed at the reciprocal of the programmed inter-reinforcement interval [(1 / *q*) - sec], this type of training does not establish such positive relationship: once sufficiently high response rates are attained, increases in responding are not followed by increases in reward rate. Assuming that responding is controlled by this response-outcome (R-O) rate correlation, it follows that ratio schedules should support higher responding than interval schedules for matched reward probabilities or rates.

Following Baum (1973; Dickinson, 1985; Dickinson & Pérez, 2017; Pérez, 2017), it could be argued that the experienced R-O rate correlation is computed by taking the number of responses and rewards experienced across a number of limited time-samples. Under this specification, participants will experience positive R-O rate correlations only if responding varies across these samples: stable or high responding narrows the rate of responding and rewards sampled and hence decreases the experienced R-O rate correlation, even in the case of ratio schedules (Dickinson & Perez, 2018).

In keeping with this argument, it is not difficult to think that the reason for the different results obtained in our study and those reported by Dawson and Dickinson (1990) concerns the different patterns of responding that may have been produced in these studies. It may well be the case that responding in Dawson and Dickinson’s (1990) was more variable across sessions than responding in the present experiment, allowing subjects to continuously re-experience positive R-O rate correlations under ratio training and bringing about higher response rates for this schedule. Of course, the most likely candidate for driving such an effect is the type of target response employed by Dawson and Dickinson. One possibility is that, in contrast with lever pressing, rats trained to pull a chain develop a pattern of responding that is more variable across and within sessions. For example, the mechanical properties (e.g. degree of hysteresis) of the microswitch from which the chains were suspended (a home-made system) may have produced sustained variation in rates of responding. By contrast, lever pressing may not produce such variability; the failure to find a difference between RR-20 and RPI-y groups in our study might simply be a consequence of lower R-O rate correlations that either failed to be detected, or else were similar between ratio and the other schedules.

We explored these possibilities by calculating the R-O rate correlations experienced by subjects early and late in training in each phase of the study. In keeping with Baum’s original idea and Dickinson and Perez’s (Dickinson & Perez, 2017; Tanaka, Balleine, & O’Doherty, 2008) modelling strategy, we divided each experimental session in 60-sec time samples, and computed a Pearson correlation coefficient between the number of responses and rewards obtained by each subject across these time-samples (see Appendix for details). Figure 2 presents the average R-O rate correlations experienced by rats in Phase 1 (left panel) and Phase 2 (right panel) of the study. Given that rats came to perform at a high and stable rate under RR-10 training, the R-O rate correlation experienced by this group converged to zero rapidly. This result confirms the widely-accepted hypothesis that the difference in responding between RR-10 and RR-20 is explained by the higher reward probability of the former. The RR-20 and RPI-y groups, in contrast, experienced positive R-O rate correlations during the first phase, but these did these did not differ either early (*t*(63) = 0.44, *p* = .66) or late (*t*(58.3) = 1.49, *p* = .14) in training.

**Fig. 2.**
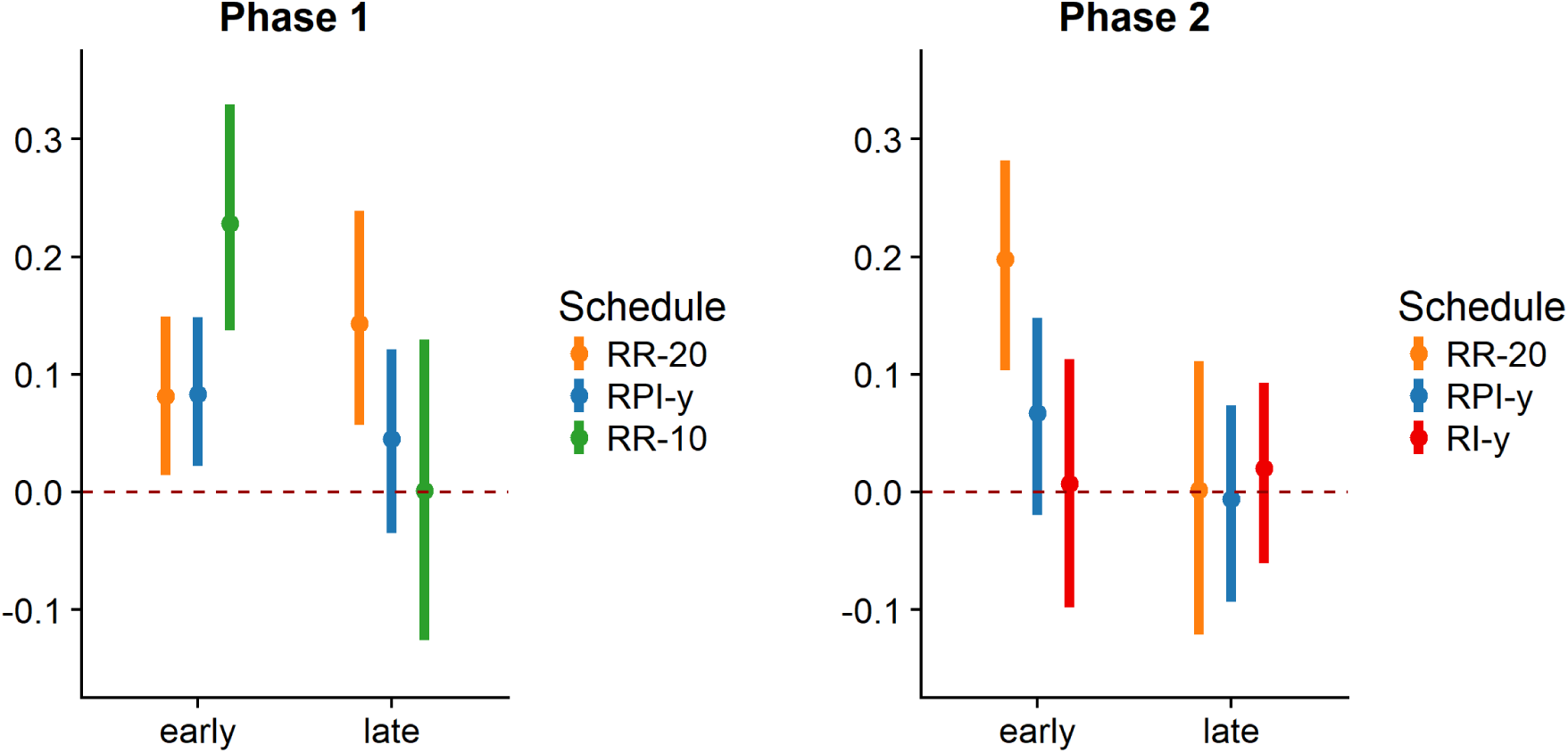
R-O rate correlations experienced by each group in Phase 1 and Phase 2 of the experiment. The values shown are mean and 95% bootstrapped confidence intervals for the first (early) and last (late) three sessions of training for each rat and group (see Appendix for details).

A clearer result was obtained in the second phase. Here, the effect of stable responding on experienced R-O rate correlations was evident in its approaching zero in all groups at the end of training (*F* (2,104) = 0.90, *p* = .41). Given the similar correlational properties of the RI-y and RPI-y, the lower performance observed in the RI-y is therefore attributable only to the differential reinforcement of long IRTs of the RI-y while the failure to detect any differences between RR-20 and RPI-y when R-O rate correlations are comparable (*t*(65.8) = 0.11, *p* = .91) is explained, again, by the matched reward rates established by the yoking procedure. These analyses confirm that the low variability in responding produced comparable R-O rate correlations between the schedules and is in agreement with the simulations presented by Dickinson and Pérez (2018).

Although these results suggest that the RR-20 schedule is not able to sustain higher lever-pressing rates than the RPI-y schedule due to its low R-O rate correlation, there is a wealth of evidence implicating this factor in instrumental conditioning. Reed (2006), for example, compared response rates on the RI-plus-linear feedback schedule (RI+) (McDowell & Wixted, 1986; Reed, 2007a, 2007b; Soto, McDowell, & Dallery, 2006) and compared this performance to that of a ratio schedule yoking reward rates. Under the RI+ specification, higher reward probabilities for long IRTs are established in conjunction with positive R-O rate correlations. Using lever pressing in rats, Reed (2006) presented evidence of an RI+ schedule sustaining similar response rates to an RR schedule for matched reward rates. However, this result was obtained only if rats were encouraged to respond at a sufficiently low rate by increasing the force required to depress the lever. When high response rates were obtained by decreasing this required force, rats responded at a higher rate in the RR than in the RI+. This result implies that these subjects were sensitive to R-O rate correlations when responses rates were low, but sensitive to the differential reinforcement of long IRTs when these were high. Consistent with our observations above, the high response rates attained by the low-force group in Reed’s study may not have allowed these rats to experience positive R-O correlations, leaving IRT-reinforcement as the main factor affecting their responding under the low force condition.

In a recent human study, Pérez and colleagues (2016) have also found evidence of R-O rate correlation mediating responding in human subjects. In a within-subject design, they trained participants under master RPI schedules with two different interval requirements. Subsequently, two RR schedules with the same reward probabilities as those obtained in the master RPI schedules were presented. Although participants responded at a similar rate in all schedules at the end of training, a final 2-min choice test showed a strong preference for the RR schedule over the RPI schedule in all groups of participants. The authors interpreted this result as a consequence of the low cost of responding during sequential training compared to the explicit opportunity cost of responding to the RPI option when the RR schedule is presented concurrently. During a choice test, participants can only control the reward rate by responding in the RR schedule; the maximum number of rewards obtainable in a limited-time trial is determined by the inter-reinforcement parameter under the RPI schedule and is therefore out of participants’ instrumental control.

Although Pérez et al. did not compare causal control judgments for the RR and RPI schedules, previous studies (Bradshaw, Freegard, & Reed, 2015; Reed, 2001) shed some light into this possibility. In these studies, participants were trained under ratio and interval schedules for matched reward rates (Exp 1.) or probabilities (Exp. 2) and asked to rate the casual effectiveness of their responses in causing the outcome. In both experiments, Reed found higher R-O causal judgments for ratio schedules compared to interval schedules - performance mirrored this result. This is, of course, in sharp contrast with the assumptions of both mechanistic and rational theories of causal learning which attribute causal judgments to some reward probability-metric such as Δ*P* = *P*(*O* | R) − *P*(*O* | *no* R) (Blanco, Matute, & Vadillo, 2013; Dickinson, Shanks, & Evenden, 1984; Hammond & Paynter, 1983; Wasserman, Chatlosh, & Neunaber, 1983) or variations of this metric (Cheng, 1997; Novick & Cheng, 2004). The most plausible explanation for Reed’s (2001) results is that participants attribute causal control in line with the correlational properties of the schedules, independently of reward rate or reward probability. Indeed, there is evidence from imaging studies suggesting a correlation between BOLD activity in the ventromedial prefrontal cortex, causal judgments of the R-O association, and the R-O rate correlation established by the schedule during training (Tanaka et al., 2008). Importantly, the result is obtained even when there was no difference in performance between ratio and interval training. Taken together, these results suggest a more complex relationship between the correlational properties of the schedules and instrumental performance. Although participants might be able to encode these properties, these will not necessarily reflect in performance unless there is an explicit cost of responding or a clear advantage in deploying this information, as in the case in which maximizing rewards is compromised by not doing it so (Pérez et al., 2016).

In spite of this evidence suggesting a role for the R-O rate correlation in instrumental learning, it is clear that a single-system theory based on this notion cannot capture the whole of the instrumental process. If, as shown above (see also Dickinson & Pérez, 2017), for example, increasing response rates are associated with lower R-O rate correlations, it follows that responding should slow down as training progresses. This lower response rate should then allow subjects to re-experience a high R-O and bring about higher response rates, contradicting the very notion of learning curve. To overcome this, Dickinson and Pérez (2017; see also Dickinson, 1985; Pérez, 2017) have recently proposed a model in which a rate-correlation system works concurrently with a reward-probability system to determine responding. Under the additional assumption that these two systems summate to control responding at any level of training, the model can account for the results observed in the present study: when the rate-correlation is equal between RR-20 and RPI-y groups, its contribution to performance should be equal; hence total responding should be determined only by the reward rate, a factor that has been explicitly controlled by yoking within triads. In addition, this dual-system model anticipates that interval training and ratio responding after extended training should be impervious to changes in the utility of the outcome, in agreement with the data reported in the instrumental conditioning literature (Daw et al., 2005; Dickinson et al., 1983; Holland, 2004).

Whereas the data reported by Pérez et al. (2016) in humans and Reed (2006) in rats suggests that, at least under some conditions, R-O rate correlation can modulate instrumental performance, the present study has clarified the ratio-RPI contrast and confirmed the predictions of IRT reinforcement theories with regard to the ratio-interval difference (Niv, 2007; Niv et al., 2007; Peele et al., 1984; Tanno & Silberberg, 2012; Wearden & Clark, 1988). If the variation of responding across sessions is not sufficient to establish different R-O rate correlations between schedules, performance on ratio schedules is comparable to that of RPI schedules; compared to RPI schedules, responding under RI schedules is slowed by the differential reinforcement of long IRTs.

## Acknowledgments

This work was funded by a UK Medical Research Council Programme Grant (G1002231) to ALM. ODP was funded by a PhD scholarship from CONICYT. ALM is the Ferreras-Willetts Fellow in Neuroscience at Downing College, Cambridge. We would like to thank Dr George Vousden for providing advice on the running of these experiments.

## Appendix

### Method for calculating R-O rate correlation

Each session of training was divided in 60-sec time samples. If the session length was not a multiple of 60, the last time-sample was discarded from the data. For each session and rat, the R-O rate correlation was calculated by taking the number of responses (*n*_*resp*_) and rewards (*n*_*rew*_) in each time-sample and calculating a Pearson correlation coefficient as 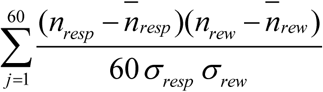, where 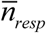 and 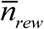 are the mean number of responses and rewards in the session, respectively, and *σ*_*resp*_ and *σ* _*rew*_ are their corresponding standard deviations. The values shown in Fig. 2 are the average of the first (*early*) or last 3 (*late*) sessions of training obtained for each rat. Since Pérez (2017)’s showed that the size of the time sample chosen does not exert an impact in the R-O rate correlations obtained when responses rates are sufficiently high, the value of 60-sec was chosen arbitrarily for this calculation.

## Footnotes

1 A similar idea been recently recognized in the computational Reinforcement Learning literature, where chunks or chain of responses are considered as *meta-actions*, or activities extended in time (Dezfouli & Balleine, 2012; Pauli, Cockburn, Pool, Pérez, & O’Doherty, 2018).

